# Pericyte contact alters endothelial cell metabolism by promoting exchange of lactate through SLC16A3

**DOI:** 10.64898/2025.12.10.693369

**Authors:** Mahdieh Safarzad, Sebastian Kempf, Roman Vuerich, Rüdiger Popp, Timo Frömel, Jiong Hu, Ingrid Fleming

## Abstract

Intimate crosstalk between endothelial cells and pericytes is fundamental for vascular development and stability, but the metabolic dimension of this interaction remains poorly defined. Using a filter-based co-culture system that mimics the shared basement membrane of the microcirculation, we performed an integrated multi-omics analysis to investigate how direct contact reprograms both cell types. We found that initial contact does not induce immediate quiescence but rather triggers a transient, low-level activation of an endothelial-to-mesenchymal transition (EndMT)-like state in endothelial cells, characterized by specific upregulation of genes involved in extracellular matrix production (e.g., collagen isoforms) and PDGFR signaling, without a full loss of endothelial identity. Concomitantly, pericytes shifted endothelial cell metabolism, increasing glycolysis and elevating intracellular pyruvate and lactate. Proteomic analysis of the contact interface revealed enrichment of solute carriers, most notably the lactate transporter SLC16A3. Functional studies demonstrated that pericytes act as a glycolytic partner, actively shuttling lactate to endothelial cells via SLC16A3. This lactate did not fuel the TCA cycle but instead served as a signaling metabolite, driving widespread alterations in the endothelial acetylome and lactylome, particularly affecting proteins involved in glycolytic metabolism. In vitro, exogenous lactate potentiated cytokine-induced EndMT and upregulated lactylation-associated genes. The physiological relevance of this lactate shuttle was confirmed in vivo, where endothelial-specific deletion of Slc16a3 in mice impaired postnatal retinal angiogenesis, leading to reduced vessel density and diminished endothelial-pericyte overlap without affecting endothelial cell proliferation. Our findings establish that vessel maturation is orchestrated by a metabolically gated phase of plasticity initiated upon first contact, wherein a targeted EndMT-like program and a pericyte-driven lactate signaling axis are integrated to coordinate vascular network assembly.

## Introduction

Endothelial cells and pericytes engage in intense, bidirectional crosstalk that regulates almost every stage of vascular biology: vessel formation, stabilization, permeability, remodeling and regression.^1^ Their communication uses secreted factors, direct cell-cell contacts and the extracellular matrix (ECM). Under normal, quiescent conditions, endothelial cells release signals that attract, support, and restrain pericytes. One of the earliest being PDGF-B, which is secreted by endothelial cells to recruit pericytes to the vessel wall and ensure that they are spaced appropriately along the capillary.^2,3^ PDGF can also stimulate metabolic communication between pericytes and endothelial cells via long thin processes referred to as tunneling nanotubes (TNTs).^4,5^ These TNTs can even transport endosomal vesicles,^6^ and organelles such lysosomes, and mitochondria.^7–9^ Once contact is established, pericytes are reported to exert a stabilizing effect and the nascent vessel releases a mutually reinforcing set of signals. For example, TGF-β promotes pericyte differentiation into a mature, contractile phenotype that helps fortify the vessel, at the same time as feeding back to suppress endothelial proliferation.^10,11^ Also, pericytes produce angiopoietin-1, which activates the Tie2 receptor on endothelial cells, tightening endothelial junctions, strengthening adhesion and suppressing the vessels tendency to sprout or leak.^12,13^ Additional stabilizing cues include sphingosine-1-phosphate (S1P),^14^ and Notch signaling.^15,16^ Alongside these soluble/cell bound signals, direct physical interactions anchor the two cell types together as they share a basement membrane characterized by peg-and-socket interdigitations.^17^ This membrane is rich in laminin, collagen IV, nidogen, and other extracellular matrix components, and contains integrins and junctional proteins as well as gap junctional plaques that facilitate the transfer of electrical signals and well as small molecules.^18^

There are conditions in which this intimate association is disrupted so that pericyte coverage decreases, which results in vessel destabilization and capillary loss. Indeed, pericyte rarefaction has been described in retinas from animals and humans with diabetes,^19^ where it contributes to non-proliferative retinopathies ^20,21^. The same phenomenon also occurs during the development of the microcirculatory dysfunction that accompanies heart failure.^22,23^ Even though endothelial cell-pericyte crosstalk maintains quiescence, stability and the proper function of capillaries, little is known about the metabolic exchange between endothelial cells and pericytes and how this can affect cell phenotype. The finding that exosomes and mitochondria can be exchanged between thetwo cell types indicates that the metabolic states of the two cell types are interdependent. Therefore, we set out to study how the cell transcriptome, proteome and metabolome are altered by cell contact and identify pathways that may be targeted to promote vessel stability.

## Results

### Co-culture reprograms endothelial cell and pericyte gene expression

We previously used a simple 2D co-culture to study endothelial cell-pericyte communication. ^9^ However, the cells tended to cluster into islands and formed tunneling nanotubes rather than the common basal lamina that characterizes the architecture of the microcirculation. For this reason, we opted for a filter based co-culture system to study endothelial cell-pericyte metabolic communication (**Fig. 1A**), that was used previously to investigate the composition of myo-endothelial junctions.^20,24^ Cells were seeded at high density and maintained in culture for 5 days to ensure that cell-cell contact could be established through the filter. RNA-sequencing (RNA-seq) was then used to determine the impact of cell-cell contact on both cell types. In pericytes (**Fig. 1B&C**), the co-culture with endothelial cells significantly decreased the expression of genes linked to synaptic/neuronal markers (*Insyn2a*, *Syt7*, *Vsnl1*), signal transduction and extracellular regulation (*Slc7a14*, Slc22A4, *Ptpru*, *Egr3*, *Adamtsl4*, *Prkg2* and *Tnfrsf* members). Among the most upregulated transcripts were genes involved in structural/extracellular matrix components (*Lmod1*, *Acan*, *Cspg4*), signaling / neuronal or vascular markers (*Notch3*, *Sorbs2*), matrix metalloproteinases and genes associated with activated or specialized cell states (*e.g. Megf6*). While changes in some of the genes indicated that pericytes cultures with endothelial cells exist in a less differentiated state (versus monoculture), other changes indicate that the cells were activated, particularly in the remodeling of membranes lipids and the extracellular matrix. Pathway enrichment analyses (Reactome and Gene Ontology biological process) identified a potential impact on pathways including ECM deposition and organization, including collagen formation, Notch signaling cytoskeletal/intermediate filament reorganization, triglyceride and cholesterol homeostasis and cell-cell junction assembly as well as inhibition of endothelial cell proliferation (**Supplementary Fig 1A&B**).

**Fig. 1.**
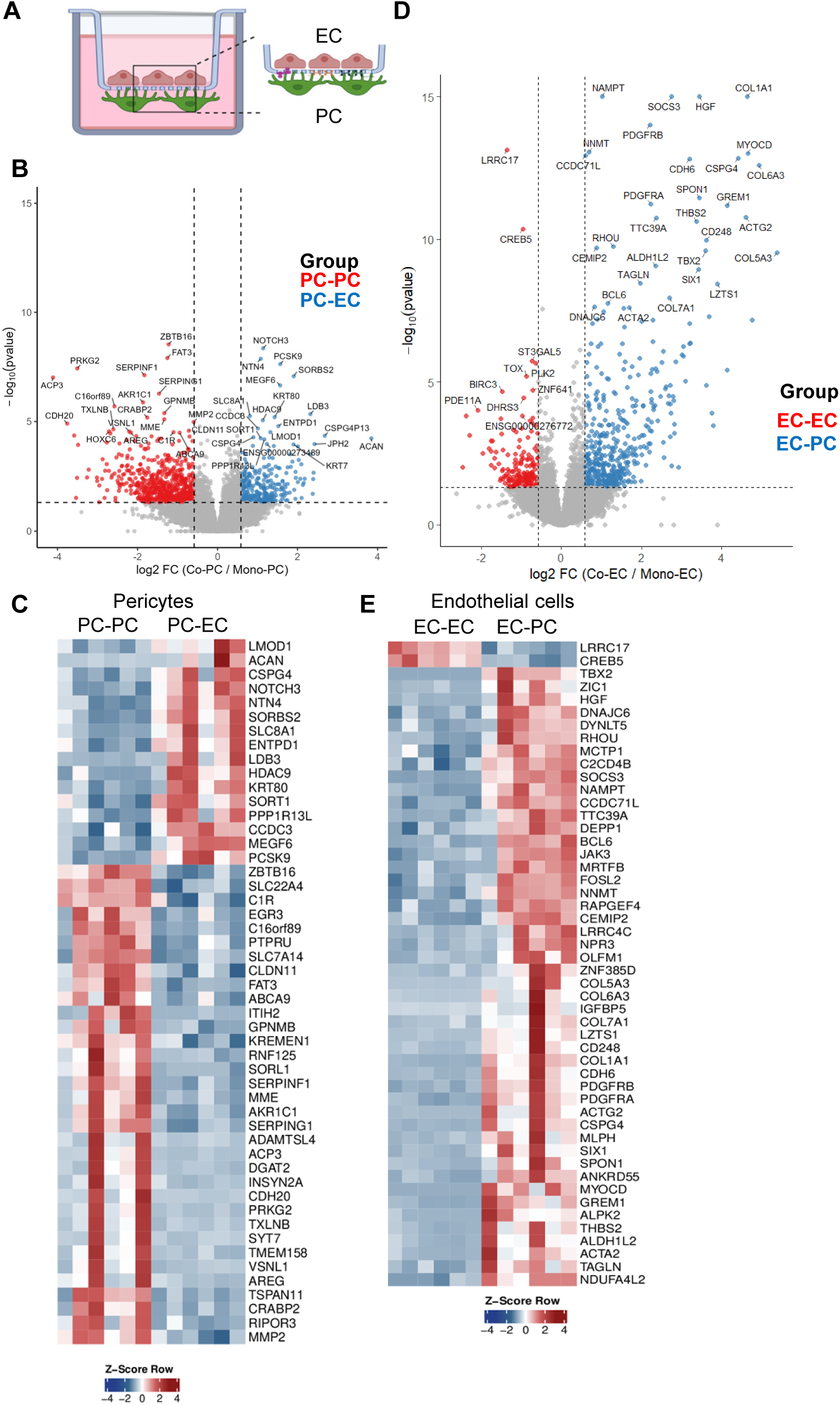
Impact of co-culture on pericyte and endothelial cell gene expression. (**A**) Filter system for endothelial cell (EC) and pericyte (PC) co-culture. **(B&C**) Volcano plot (A) and heatmap (B) showing the pericyte genes highly expressed in monoculture (PC-PC) versus co-culture with endothelial cells (PC-EC); n=6 independent cell batches. (**D&E**) Volcano plot (C) and heatmap (D) showing the endothelial cell genes highly expressed in monoculture (EC-EC) versus co-culture with pericytes (EC-PC); n=6 independent cell batches.

In endothelial cells, co-culture with pericytes altered gene expression consistent with the activation of a program to generate ECM and some of the changes in transcripts indicated a discrete activation of mesenchymal genes, indicating a partial endothelial to mesenchymal transition (EndMT) state (**Fig. 1D&E**). Among the genes most upregulated in endothelial cells in contact with pericytes were *Hgf*, *Pdgfra*, *Pdgfrb*, *Socs3*, *Jak3*, *Nampt*, *Nnmt*, *Cemip2*, as well as *Col5a3*, *Col6a3*, *Col7a1*, *Col1a1*, *Mlph* and *Spon1*. Particularly indicative of EndMT were the changes in *Grem1*, *Actg2*, *Acta2*, *Tagln*, *Myocd*, *Cd248* and *Thbs2*. Pathway enrichment analyses identified collagen biosynthesis, modification, and ECM organization as the pathways most upregulated in endothelial cells in co-culture. Among the pathways depressed by pericytes were TNF signaling, necrosis, necroptosis, and NF-κB signaling (**Supplementary Fig 1C&D**).

### Impact of pericytes on endothelial cell metabolism

RNA-seq, although it provides useful information, does not always represent changes in protein expression. Therefore, we performed proteomics to detect the impact of co-culture on the expression of enzymes involved in metabolism. Because we previously observed mitochondrial transfer from pericytes to endothelial cells and pericytes had a more pronounced effect on endothelial cell gene expression we concentrated on the endothelial cell proteome. This approach revealed a significant impact of pericytes on the endothelial cell expression of enzymes involved in glycolysis, the pentose phosphate pathway and the TCA cycle as well as amino acid and fatty acid metabolism (**Fig. 2A**). Targeted metabolomics focusing on selected intermediate of glycolysis and the TCA cycle revealed that co-culture with pericytes induced a metabolic shift in endothelial cells (**Fig. 2B**). Contact with pericytes increased endothelial cell glucose 6-phosphate, pyruvate and lactate levels, indicating increased glycolysis. Levels of endothelial cell ribulose-5-phosphate were decreased by pericytes indicating redirection of glucose away from the pentose phosphate pathway. Despite the high levels of pyruvate and lactate, isocitrate content was comparable in endothelial cells in monoculture and in co-culture with pericytes. Conversion of glutamine to α-ketoglutarate was increased by pericyte contact but this did not appear to feed into the TCA cycle as succinate levels were also comparable in both groups.

**Fig. 2.**
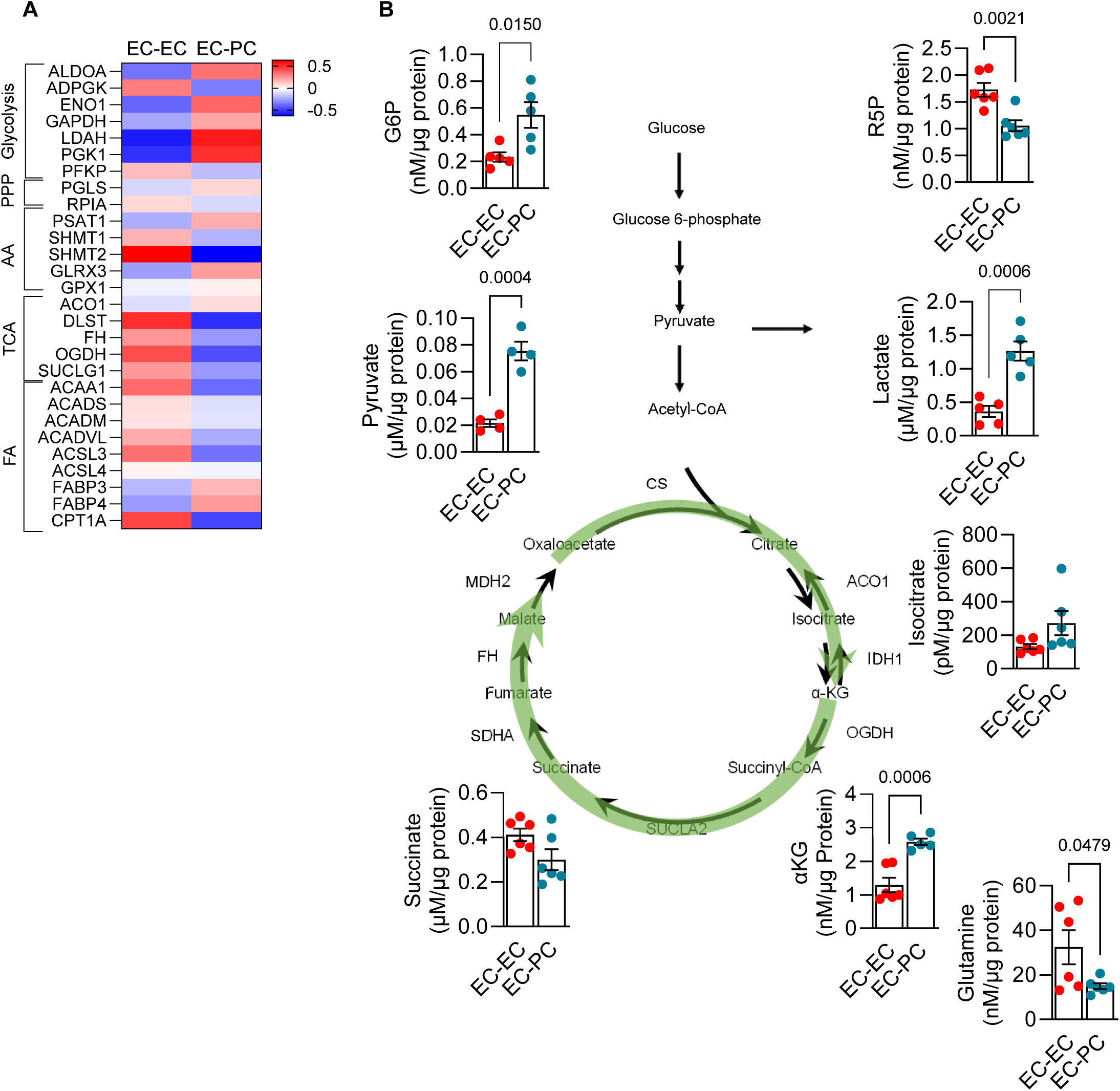
Impact of pericytes on endothelial cell metabolism. (**A**) Heat map showing the relative expression of key metabolic enzymes in endothelial cells in monoculture (EC-EC) or in co-culture with pericytes (EC-PC). Enzymes are grouped by major metabolic pathways, including glycolysis, the pentose phosphate pathway (PPP), amino acid (AA) metabolism, the tricarboxylic acid (TCA) cycle, and fatty acid (FA) metabolism. Color scale indicates normalized expression values. (**B**) Metabolite profiling of glycolysis and TCA cycle intermediates in endothelial cells in monoculture (EC-EC) versus co-culture with pericytes (EC-PC); n=4-6 independent cell batches. (Student’s t test).

Coculture with pericytes had a more global impact on the endothelial cell proteome (**Fig. 3A&B**, **Supplementary Fig. 2**) as co-culture decreased the endothelial cell expression of proteins involved in fatty-acid β-oxidation / lipid metabolism, such as short/branched-chain acyl-CoA dehydrogenase (ACADSB) and ceramide synthase 2 (CERS2). Several metabolic enzymes were increased by pericyte contact including phosphoglycerate kinase 1 (PGK1) phosphoglucomutase-like enzyme (PGM3) which links glucose-6-phosphate handling and the hexosamine pathway, LETM1 – mitochondrial inner-membrane protein involved in ion homeostasis that indirectly affects TCA/respiration, and NAMPT a key NAD⁺ salvage enzyme. Pericyte contact also increased JunB levels which although it is upregulated by VEGF and other pro-angiogenic signals it also plays a role in vascular maturation processes.^25,26^ While some of the changes observed e.g. the suppression of CDK1, would fit well with the well-established concept that pericytes promote endothelial cell quiescence there was also clear evidence of endothelial cell activation that fit well with the RNA-seq data.

**Fig. 3.**
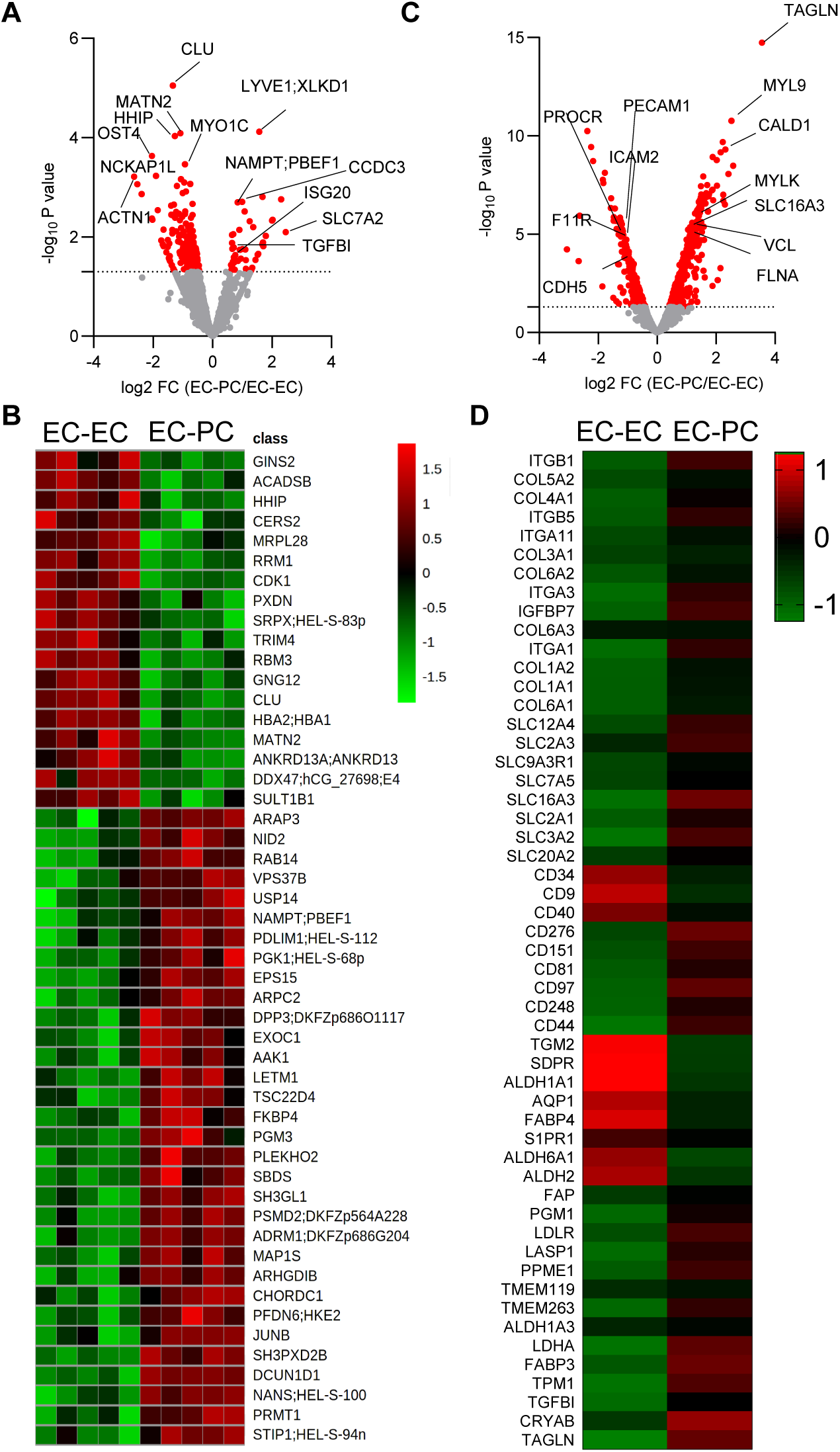
Impact of pericytes on the endothelial cell proteome. (**A**) Volcano plot displaying significantly altered proteins in endothelial cells in monoculture (EC-EC) versus coculture with pericytes (EC-PC). Labeled points represent the top significantly regulated targets. (**B**) Heatmap (Z-score–scaled abundance values) of the 40 most differentially expressed proteins as in A; n= 5 independent cell batches. (**C**) Volcano plot showing differential expression of proteins in filter pores from endothelial cells in monoculture (EC-EC) versus co-culture with pericytes (EC-PC); n= 6 independent cell batches. (**D**) Heatmap (Z-score normalized filter protein expression) as in C. Z-score normalization was applied per protein to highlight relative expression differences.

To assess potential mechanisms involved in endothelial cell-pericyte communication we repeated our analyses using proteins recovered from the filter pore that were enriched in endothelial cell-pericyte junctions (**Fig. 3C&D**). While CD31 (PECAM1) and CDH5 were enriched in filters from endothelial cell cultures, pericyte-endothelial cell contacts were enriched in numerous collagen and integrin isoforms, as well as several members of the solute carrier (SLC) family e.g. the ion transporters SLC12A4 (potassium and chloride) and SLC20A2 (phosphate), the glucose transporters SLC2A1 (GLUT-1) and SLC2A3 (GLUT3), L-type amino acid transporters SLC7A5 and SLC3A2 and the monocarboxylate transporter SLC16A3.

### Lactate shuttles via SLC16A3 between pericytes and endothelial cells

SLC16A3 (also known as MCT4) was also among the most differentially regulated proteins and is a monocarboxylate transporters that can transport lactate and pyruvate as well as ketone bodies such as D-β-hydroxybutyrate and acetoacetate. Whether the net result is influx or efflux depends on the concentration of the substrates as well as pH gradients.^27^ Given the marked upregulation of SLC16A3 in filters from the co-culture, together with the observation that pericytes induced the accumulation of lactate and pyruvate in endothelial cells, we determined whether lactate could shuttle between the two cell types. Western blotting confirmed the enrichment of SLC16A3 in filters from co-cultured endothelial cells and pericytes (**Fig. 4A**). To determine whether endothelial cell pericyte junctions in vivo are also enriched in the transporter we assessed its expression the cardiac microcirculation of wild-type mice. SLC16A3 was located at sites of endothelial cell-pericyte contact peaking with the signals for CD31 and NG2 (**Fig. 4B**). Next, a short pulse of ^13^C glucose was used to assess flux to lactate. In monocultures of endothelial cells, this resulted the appearance of ^13^C-labelled glucose-6-phosphate, pyruvate and lactate (**Fig. 4C**). In the presence of pericytes, however, flux to lactate was significantly elevated and was higher than could have fluxed from pyruvate. To determine whether this excess endothelial cell lactate could shuttle from pericytes, targeted metabolomics was used to assess lactate levels in the absence and presence of the lactate transporter inhibitor, syrosingopine. As before contact with pericytes increased endothelial cell lactate but this phenomenon was not observed in inhibitor treated cells (**Fig. 4D**). The opposite was observed in pericytes i.e. coculture with pericytes decreased lactate levels in a syrosingopine-sensitive manner (**Fig. 4E**). Because only ^13^C lactate levels were affected, we hypothesized that ^13^C lactate was actively imported into endothelial cells in co-culture with pericytes via SLC16A3.

**Fig. 4.**
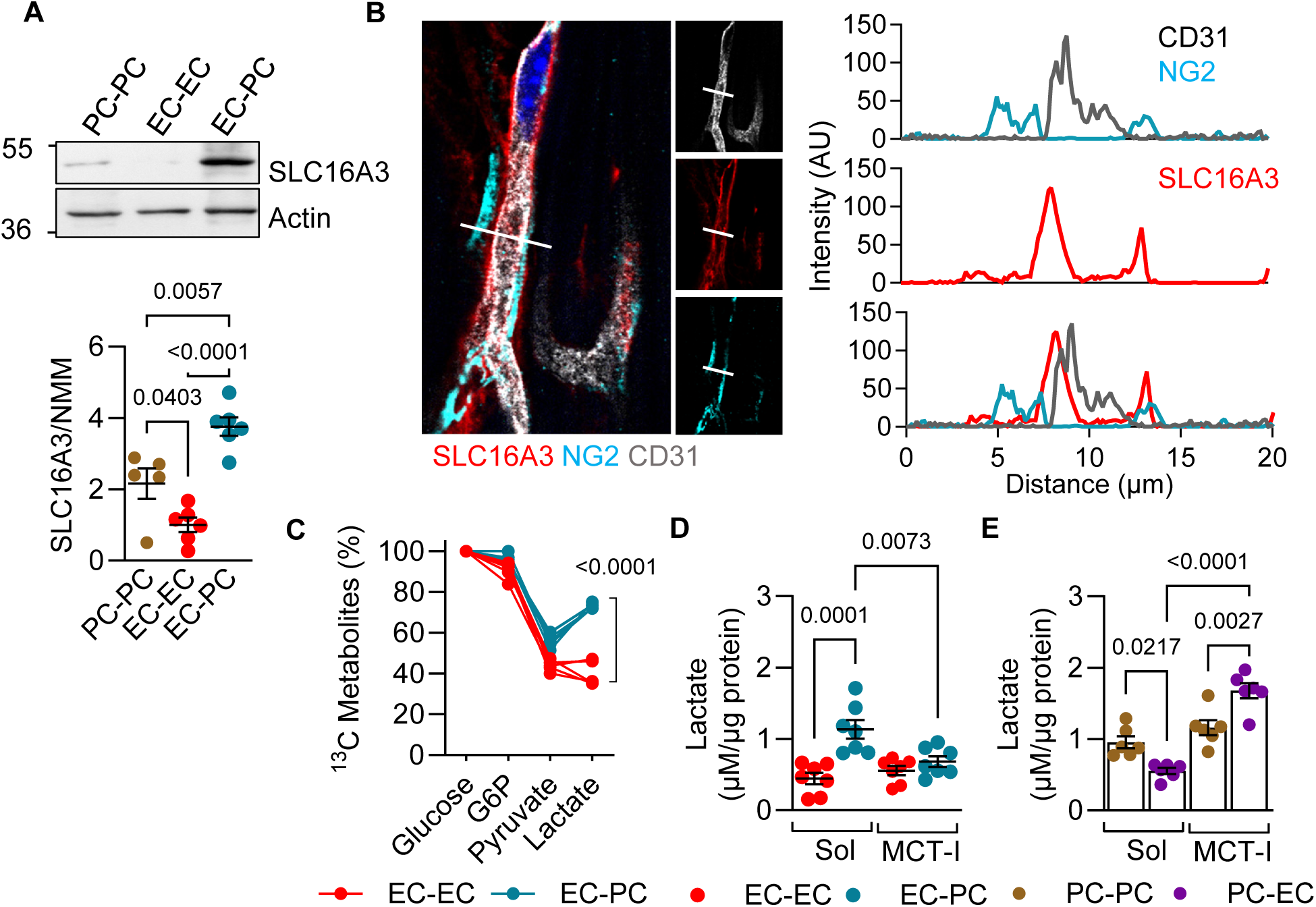
SLC16A3 in endothelial cell-pericyte junctions transfers lactate from pericytes to endothelial cells. (**A**) SLC16A3 expression in pericytes (PC-PC) and endothelial cells (EC-EC) in monoculture versus endothelial cell-pericyte co-cultures (EC-PC); n=5 independent cell batches. One way ANOVA and Tukey’s multiple comparisons test. (**B**) SLC16A3 in endothelial cell (CD31) pericyte (NG2) junctions in the adult murine heart. The image is representative of 3 additional experiments. (**C**) Flux of ^13^C-glucose to lactate in endothelial cells in monoculture or co-culture with pericytes; n=6 independent cell batches. Two way ANOVA and Sidak’s multiple comparisons test. (**D**) Lactate concentrations in endothelial cells in monoculture (EC-EC) versus co culture with pericytes (EC-PC) in the presence of solvent (Sol) or syrosingopine (MCT-I): n=5 independent cell batches. Two way ANOVA and Tukey’s multiple comparisons test. (**E**) Lactate concentrations in pericytes in monoculture (PC-PC) versus co culture with endothelial cells (PC-EC) in the presence of solvent (Sol) or syrosingopine (MCT-I) n=5 independent cell batches. Two way ANOVA and Tukey’s multiple comparisons test.

### Pericyte coculture remodels global endothelial acetylation and lactylation

Once transferred, lactate could enter the TCA cycle to generate acetyl-CoA, this could result in increased protein/histone acetylation (Kac). It could also result in another recently described modification i.e. lysine lactylation (Kla).^28,29^ Using pan-Kac antibodies we detected particularly high signals in endothelial cells that were decreased after contact with pericytes (**Fig. 5A**). Looking at Kac in more detail using mass spectrometry revealed 52 differentially (increased and decreased) acetylated lysine’s in endothelial cell proteins in mono-versus co-culture (**Fig 5B**), in addition to 48 modified histone residues (**Fig. 5C**). Modifications were largely linked to metabolism, in particular pyruvate, glucose and lactate metabolism (**Fig. 5D**). The lactylation of endothelial cell proteins and histones was also altered by co-culture with pericytes (**Fig. 5E-G**) and Panther enrichment analysis again identified glycolysis as the pathway most affected (**Fig. 5H**). These post-translational modifications provide an additional regulatory layer linking metabolic changes to protein function.

**Fig. 5.**
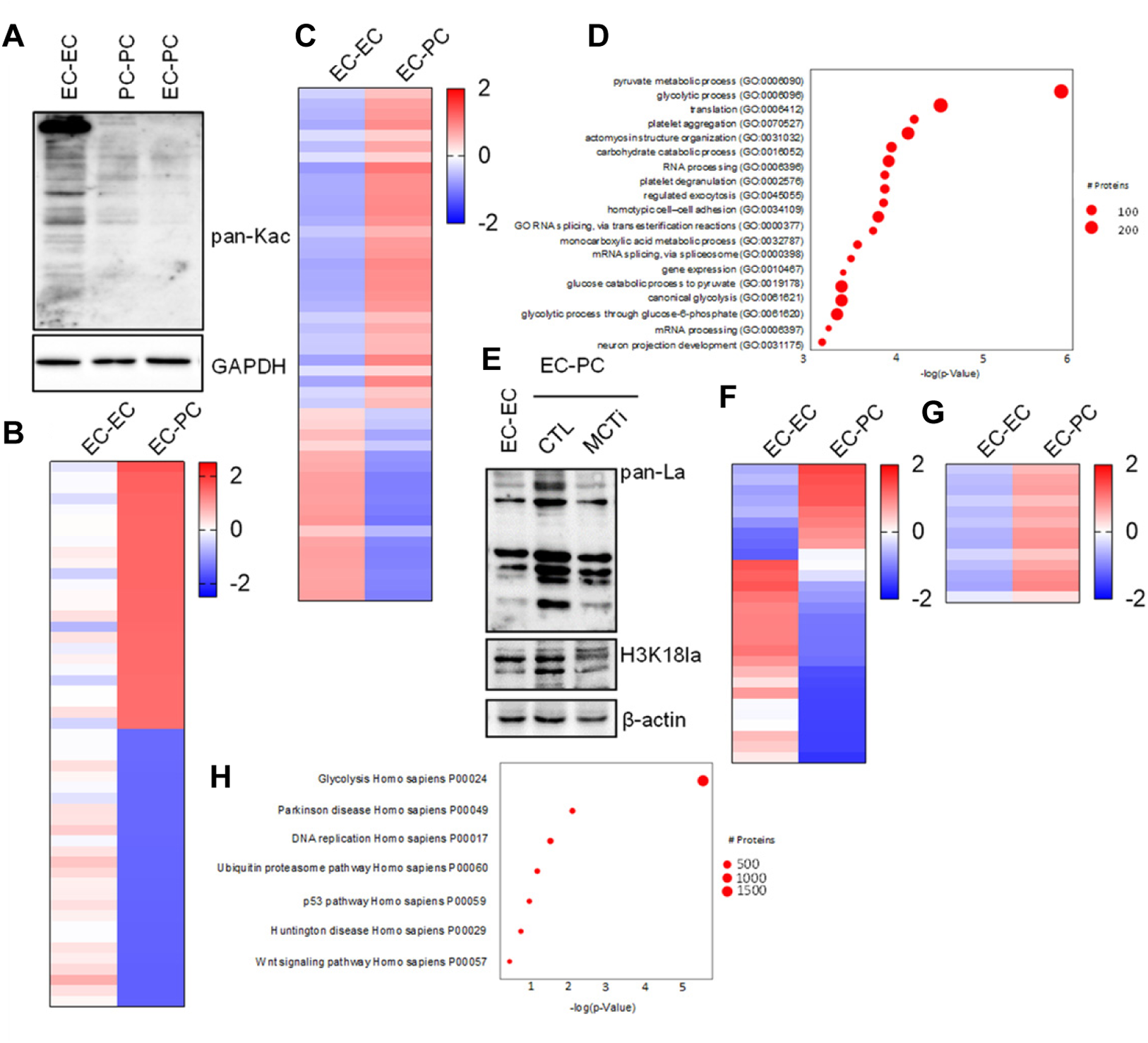
Impact of pericytes on lysine acetylation and lactylation in endothelial cells. (**A**) Global lysine acetylation (pan-Kac) in endothelial cells and pericytes in monoculture (EC-EC) versus endothelial cells in coculture with pericytes (EC-PC). (**B&C**) Top 50 significantly regulated protein Kac sites in endothelial cell proteins (B) and histones (C); n=3 independent cell batches. (**D**) Pathway enrichment analysis (GO Biological process) of proteins exhibiting significantly altered acetylation in endothelial cells. (**E**) Global lysine lactylation (pan-Kla) of proteins in H3K18 in endothelial cells in mono-versus co-culture. (**F&G**) Top 50 significantly regulated lactylation sites in endothelial cell proteins (F) and histones (G); n=3 independent cell batches. (**H**) Pathway enrichment analysis (Panther) of proteins exhibiting significantly altered lactylation in endothelial cells.

### Lactylation and EndMT

The multi-omics approach identified a number of pericyte-induced changes in endothelial cells including the activation of an EndMT-like program. Therefore, we assessed the impact of EndMT induction using IL-1β and TGF-β2 on the expression of lactylation-associated genes. The cytokine cocktail induced the expression of the EndMT markers TAGLN and αSMA, a phenomenon that was markedly enhanced in the presence of extracellular lactate (**Fig. 6A**). In parallel, bulk RNA sequencing confirmed the upregulation of additional mesenchymal markers e.g. *Cnn2* and *Fn1* (**Fig. 6B**). The expression of SLC16A1 and SLC16A3 was also increased in cells undergoing EndMT, as were enzymes involved in lactylation i.e., *Glo1*^30^ and *Aars1*.^31^.

**Fig. 6.**
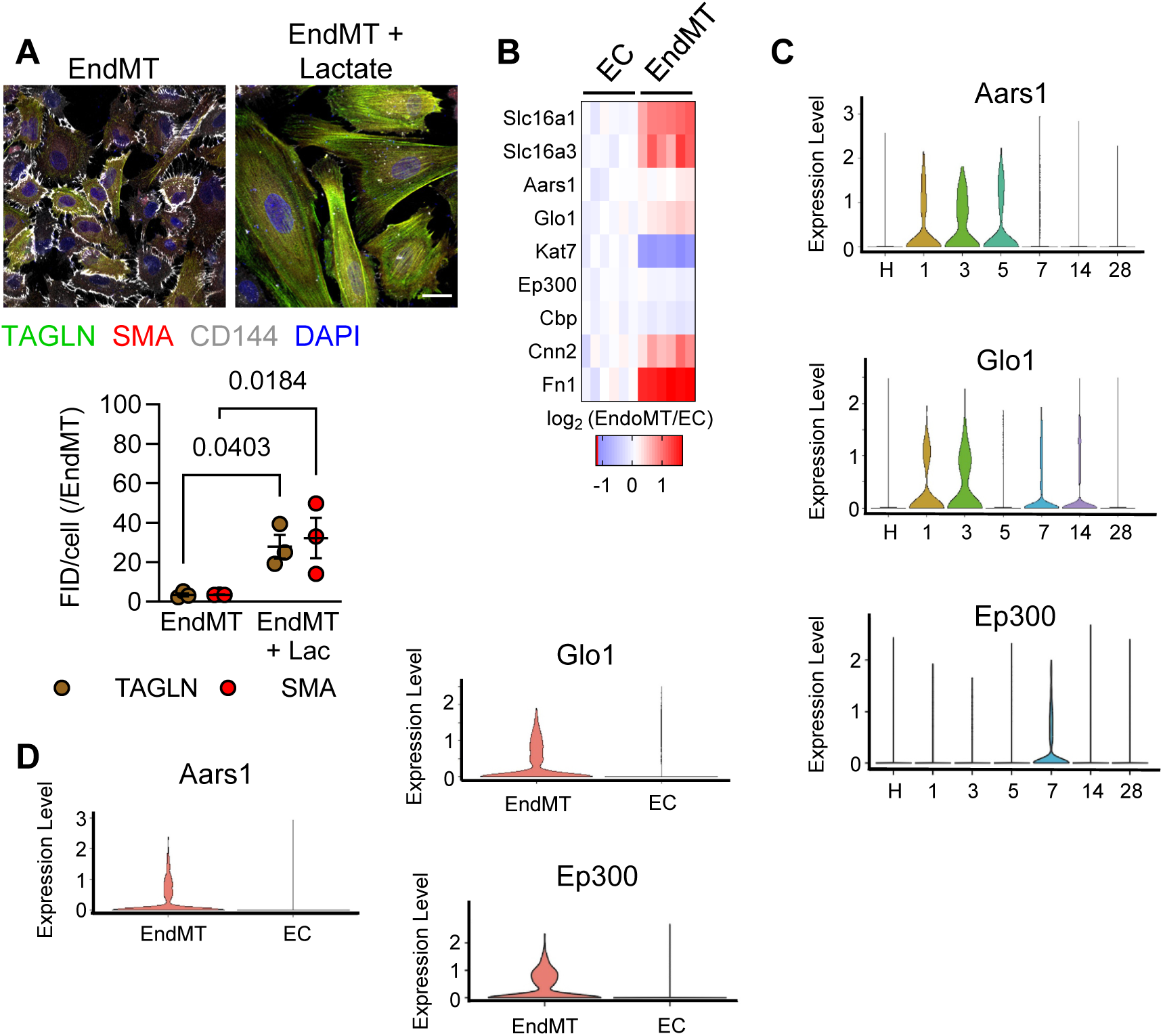
Expression of lactylation-associated genes is upregulated in ECs during EndoMT. (**A**) Confocal images of endothelial cells treated with IL1β and TGFβ2 to stimulate EndMT in the absence and presence of lactate. The graph shows mean fluorescence integrated density (FID) of TAGLN and aSMA ; n= 3 independent cell batches. Bar = 20 µm (**B**) Impact of EndMT induction on the mesenchymal markers *Cnn2* and *Fn1* and lactylation-associated genes; n=6 independent cell batches. (**C&D**) Violin plots (scRNAseq) comparing the expression of lactylation-associated genes *Glo1*, *Aars*, *Ep300* in cardiac endothelial cells at different times during EndMT after myocardial infarction in mice (C). Expression of lactylation-associated genes in naïve endothelial cells versus cells undergoing EndMT (D); Data represent 1 mouse per timepoint. Endothelial cells we selected as *Cdh5+ CD31+* cells; EndMT as *Cdh5+ CD31+ Fn1+ Cnn2+* cells.

Interrogation of a previously published scRNA-seq dataset^32^ from mouse hearts before and after MI revealed a similar, temporally restricted regulation of the same lactylation-associated genes. In Cdh5+ Cd31+ endothelial cells, *Glo1*, *Aars1* and *Ep300* expression increased during the first week after myocardial infarction i.e., when endothelial cells undergo to EndMT. By day 28 the expression these genes had returned to levels detected in naïve cells (**Fig. 6C**). The population of endothelial cells undergoing EndMT contained more cells expressing these lactylation-related genes than the cluster of naïve endothelial cells (**Fig. 6D**). Together, these data indicate that the lactate transport and lactylation machinery is selectively engaged during EndMT both in vitro and in vivo, supporting the concept that a dynamic lactylation axis contributes to the transient EndMT-like state of cardiac endothelial cells.

### Endothelial SLC16A3 deletion impairs vascular network development in the retina

To assess the impact of SLC16A3 on endothelial cell-pericyte communication Slc16a3^fl/fl^ mice were crossed with tamoxifen-inducible Cdh5-CreERT2 mice,^33^ to generate Slc16a3^iΔEC^ mice that lacked SLC16A3 in endothelial cells (**Supplementary Fig. 3A**). Retinal angiogenesis was then assessed on postnatal day (P) 6 in retains from pups that received tamoxifen daily from P1-4 (**Fig. 7A**). The endothelial cell-specific deletion of SLC16A3 did not have a significant impact on endothelial cell outgrowth from the optic nerve to the periphery as coverage was comparable in Slc16a3^fl/fl^ and Slc16a3^iΔEC^ littermates (**Fig. 7B**). However, while the coverage of nascent arteries with NG2+ pericytes was evident in retinas from the control group, coverage was clearly reduced in Slc16a3^iΔEC^ mice (**Fig. 7C**). There was also a decrease in pericyte coverage of capillaries as well as the overlap of NG2 with CD31 in Slc16a3^iΔEC^ retinas (**Fig. 7D**). Overall, the density of the vascular network in the SLC16A3^iΔEC^ retinas was much less than in control littermates (**Fig. 7E&F**, **Supplementary Fig. 3B**). Consistent with the fact that the radial outgrowth of endothelial cells was comparable between the two genotypes, endothelial cell deletion of SLC16A3 did not affect endothelial cell proliferation (**Fig. 8A**). There was also a decrease in pericyte coverage towards the angiogenic front and SLC16A3 deletion in endothelial cells impacted on the number and length of filopodia on tip cells (**Fig. 8B-D**).

**Fig. 7.**
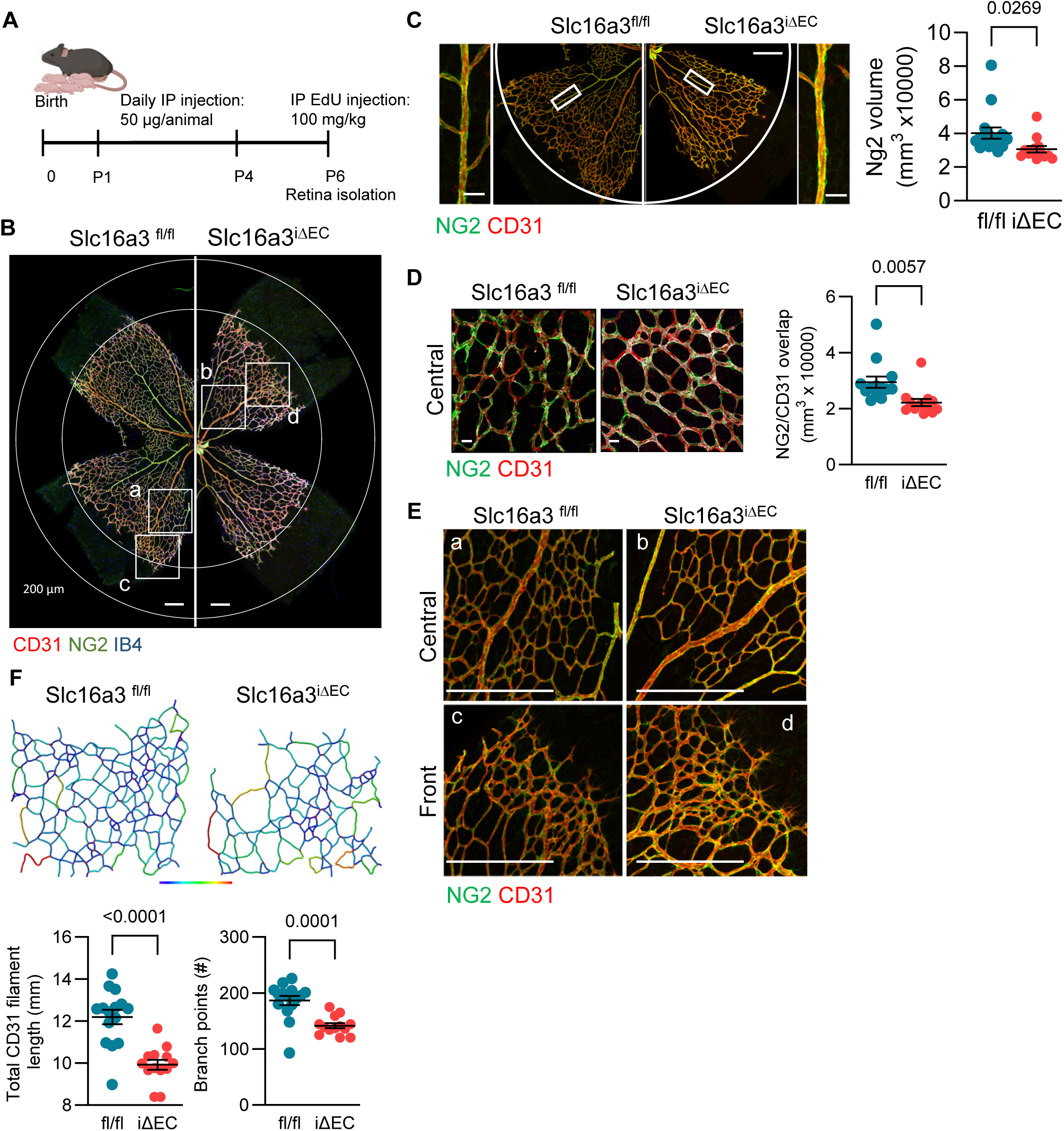
Impact of endothelial cell-specific SLC16A3 deletion on retinal angiogenesis on P6. (**A**) Scheme of the experimental protocol. (**B**) Representative whole-mount images of retinas from Slc16a3^fl/fl^ and Slc16a3^iΔEC^ mice on P6. Bar = 200 µm. (**C**) Pericyte (NG2+) coverage of nascent arteries from the same image as in A. Panels at the side are magnifications of the areas highlighted by white boxes; n=6-7 mice, 2 retinas / mouse. Bar = 300 µm whole-mount, 200 µm side panels. (**D**) Pericyte (NG2+) coverage of capillaries in the central region; n=6-7 mice, 2 retinas / mouse. Bar = 20 µm. (**E**) Vessel network in the central and front sections of the retina as indicate din B. Bar = 300 µm. (**F**) 3D segmentation of the vascular network with segment lengths visualized by a color gradient spanning the range of 2 – 100 µm (blue to red); n= n=6-7 mice, 2 retinas / mouse C, D & F: Student’s t test.

**Fig. 8.**
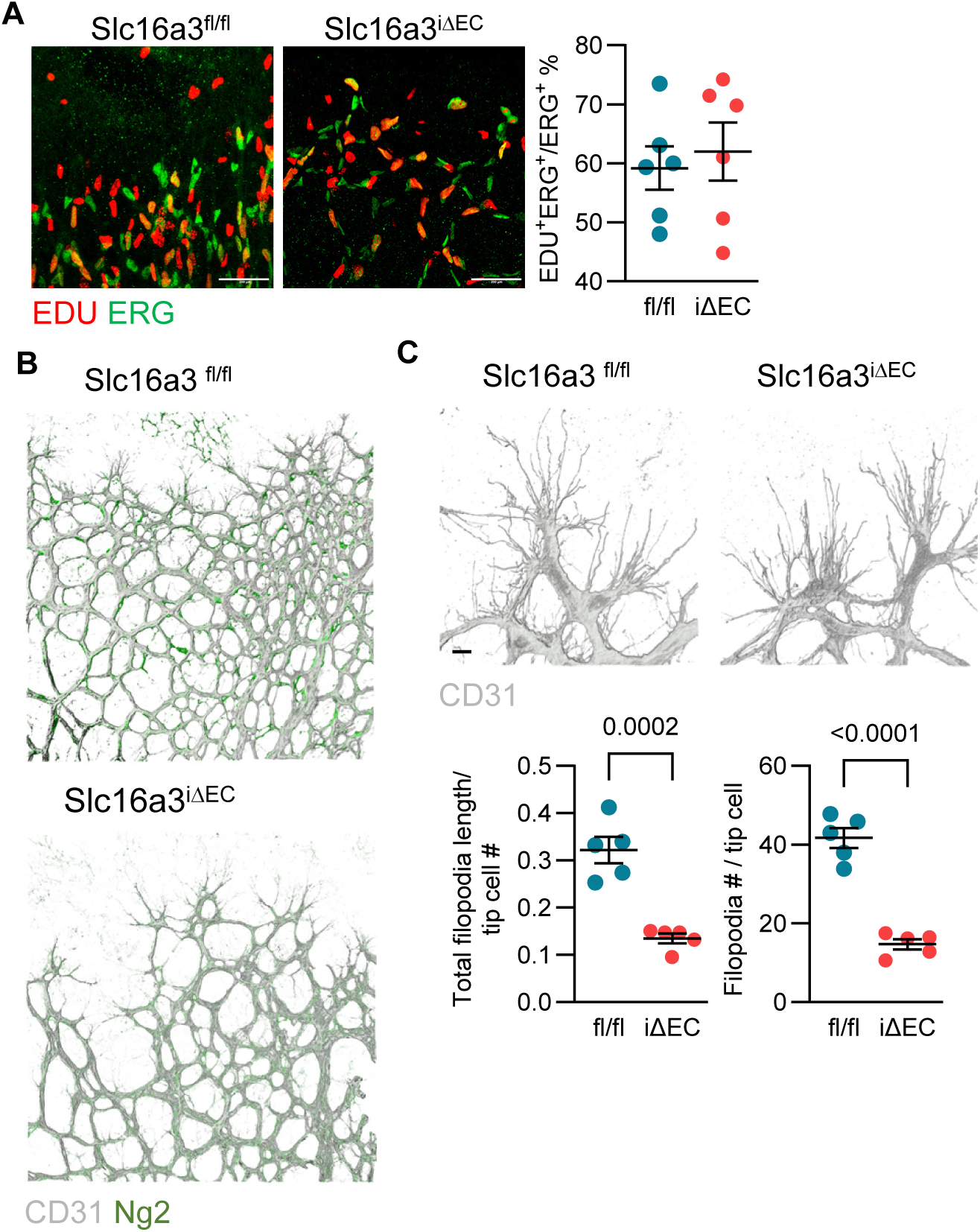
Impact of endothelial cell-specific SLC16A3 deletion on retinal angiogenesis on P6. (**A**) Representative whole-mount images of retinas from Slc16a3^fl/fl^ and Slc16a3^iΔEC^ mice on P6 showing endothelial cell proliferation (EDU+ ERG+); n=6 mice/group. Bar = 300 µm. (**B**) Pericyte (NG2+) coverage of tip cells at the angiogenic front. (**C**) Tip cells and filipodia in P6 retinas from Slc16a3^fl/fl^ and Slc16a3^iΔEC^ mice; n= 5 mice/group. Bar = 10 µm. A&D: Student’s t test.

## Discussion

The results of this investigation revealed that contact between endothelial cells and pericytes exerts reciprocal effects on both cell types to alter activation states, gene and protein expression, as well as metabolism. Overall, the impact of pericytes on endothelial cells was greater than that of endothelial cells on pericytes, which is the reason why the study concentrated only on changes in the endothelium. Some of the most prominent changes were in the production of collagen and other ECM components needed to form a stable matrix/basal lamina. The early phase of contact was, however, not characterized by a classical endothelial cell quiescence but by the initiation of an EndMT-like state. Contact with pericytes also altered endothelial cell metabolism and increased endothelial cell pyruvate and lactate levels that were at least partly due to their transfer via SLC16A3. This altered the lactylation of endothelial cell proteins and the endothelial cell specific deletion of SLC16A3 altered the development of the vascular network in the mouse retina. Although the loss of the lactate transporter did not have a substantial effect on endothelial cell proliferation at the angiogenic front, vessel density was clearly reduced as was the endothelial cell-pericyte overlap.

Pericytes can drive an EndMT by secreting active TGF-β, PDGF, and other cytokines.^1^ Indeed, the gene expression changes detected in endothelial cells that indicated the activation of TGF-β/Smad and PDGF/PDGFR signalling to increase the into expression of genes like PDGFRB that can affect migration,^34^ or cytoskeletal changes via RHOU.^35^ However, rather than co-culture inducing full-blown, EndMT we observed a low-*level* activation of select EndMT-associated pathways. For example, although genes such as *Tagln and Grem1*, which is a key regulator of EndMT,^36,37^ were induced in endothelial cells by pericyte contact, there was no detectable increase in Snail or Twist and VE-cadherin levels were unaltered. It is therefore tempting to propose that development/repair are associated with an "EndMT-like" state that is transient and self-limiting and that once stable contact is made and ECM deposited, stabilizing signals rapidly dominate and shut down the plastic program to drive both cells toward quiescence.

While the role of growth factors in and growth factor receptors in mediating endothelial cell-pericyte crosstalk has been studied in detail,^38,39^ much less is known about direct metabolic communication. That this occurs has been highlighted in several studies that reported even the transfer of mitochondria from pericyte to pericyte as well as from pericyte to endothelial cell.^9,40,41^ We therefore used RNA-seq and proteomics to assess the impact of pericytes on endothelial cell metabolism. Several pathways were affected by co-culture but the largest effects were on glycolysis, which was increased by pericyte contact, and the TCA cycle several components of which were decreased. At the level of function, pericytes promoted the accumulation of pyruvate and lactate in endothelial cells without increasing activity of the TCA cycle as isocitrate and succinate levels were comparable in endothelial cells in monoculture versus co-culture. Pericytes also increased glutamine use by endothelial cells and increased α-ketoglutarate production. As this occurred despite the accumulation of pyruvate and lactate it seems likely that glutamine was used for non-TCA-related functions, such as the synthesis of essential biomolecules (lipids, amino acids) and epigenetic methylation reactions.^42^

As endothelial cells and pericytes share a common basal lamina we assessed the composition of endothelial cell-pericyte junctions in the filter of the co-culture system to ascertain how communication could occur. This identified the enrichment of a series of proteins belonging to solute carrier family of proteins, including the moncarboxylate transporter, SLC16A3. Using the combination of targeted metabolomics and metabolic flux experiments, it was possible to demonstrate that the pericyte-induced accumulation of endothelial cell lactate depended on the activity of the transporter. This resembles the “lactate shuttle” concept first reported in recognized in muscle and neural tissue, where lactate exported from glycolytic support cells (e.g. astrocytes) is taken up by oxidative cells (e.g. neurons) via high-affinity transporters.^43,44^ Maximum glucose transport capacity is higher in pericytes than in endothelial cells, which is likely to result in increased lactate production as well as glucose oxidation,^45^ so that in our system pericytes act as the glycolytic partner delivering lactate to endothelial cells. Ours is not the first report of a role for lactate in endothelial cell-pericyte crosstalk. Indeed, the transfer of lactate from endothelial cells to pericytes was proposed to support energy generation and amino acid biosynthesis. Without endothelium-derived lactate which was achieved by preventing GLUT1-mediated glucose uptake pericyte-endothelial cell contacts were destabilized and barrier function was compromised.^46^ The reasons for the rather contradictory conclusions are probably related to the experimental model studied as while lactate transfer from the endothelium was reported to occur on first contact with pericytes, we used a longer term culture system where the cells were in culture for 5 days and had established an ECM.

The next question was what the consequence of lactate accumulation in endothelial cells could be, especially as pericyte co-culture did not increase TCA cycle activity and glutamine use. Given the parallel accumulation of α-ketoglutarate we focussed on pericyte-induced changes in protein and histone acetylation and lactylation. Pericytes had a marked effect on acetylation with differential effects on different targets. Overall, the changes in acetylation had a large impact on pyruvate metabolism and glycolysis. Lactylation was also clearly altered in endothelial cells following contact with pericytes and although a global increase in lysine lactylation was detected by Western blotting, different subsets of proteins were lactylated /de-lactylated in the co-culture situation. Not a lot is known about protein lactylation in endothelial cells but increased lactate levels have been reported to trigger EndMT by lactating proteins such as Snai1^47–49^ and Twist,^50^ among others. Certainly, lactate enhanced EndMT induction by TGF-β and IL-1β in vitro and increased expression of SLC16A3 and the lactylation “writers” Aars1 and Glo 1. Interrogation of a publicly available dataset,^32^ also revealed the transient upregulation of the same genes in native cardiac endothelial cells undergoing EndMT after myocardial infarction. To determine the consequences of SLC16A3-mediated lactate shuttling in vivo we generated mice that lacked the transporter specifically in endothelial cells. The deletion of the lactate transporter had no consistent effect on endothelial cell proliferation in the retina but had a marked impact on vessel maturation, resulting in vessel rarefaction and decreased endothelial cell-pericyte overlap.

Taken together, this study reveals that initial contact between endothelial cells and pericytes initiates a transient, metabolically-driven activation state essential for vessel maturation, rather than immediate quiescence. We observed a low-level, partial activation of EndMT-associated pathways marked by specific gene expression changes without full commitment that facilitates integration and matrix deposition. Crucially, we identified direct metabolic communication as a core mechanism. Pericytes act as a glycolytic partner, transferring lactate to endothelial cells via the SLC16A3 transporter. This lactate functions not as a fuel but as a signaling metabolite, reprogramming the endothelial epigenome through widespread changes in protein lactylation and acetylation. Disrupting this lactate shuttle in vivo by endothelial-specific deletion of Slc16a3 impaired retinal vascular development, reducing vessel density and endothelial-pericyte overlap without affecting proliferation. Our findings have broader implications for understanding tissue development, regeneration, and diseases of vascular destabilization.

## Materials and methods

### Animals

Slc16a3 floxed mice (C57BL/6Smoc-*Slc16a3^em1(flox)Smoc^*) were generated by Shanghai Model Organisms (Shanghai, China), by inserting loxP sites flanking exons 3-6. Mice were and crossed with tamoxifen-inducible Cdh5-CreERT2 mice^33^ to create mice with the tamoxifen-inducible, endothelial-specific deletion of Slc16a3 (Slc16a3^iΔEC^ mice). To induce endothelial-specific recombination, neonatal mice received intraperitoneal tamoxifen (Cay13258-10, Cayman Chemical, Ann Arbor, Michigan, USA) once daily from P 1-4 (50 µg per pup i.e. 25 µL of a 2 mg/mL solution).

Mice were maintained under circumstances consistent with the Guide for the Care and Use of Laboratory Animals published by the U. S. National Institutes of Health (NIH publication no. 85-23). All experiments were approved by the governmental authorities (Regierungspräsidium Darmstadt: FU2106). Age- and strain-matched animals of both sexes (usually littermates) were used throughout. For the isolation of organs, mice were sacrificed using 4% isoflurane in air and subsequent exsanguination.

### Retinal angiogenesis

Retinal vascular morphology and endothelial proliferation were assessed on P6. Four hours prior to sacrifice pups received an intraperitoneal injection of EdU (5-ethynyl-2′-deoxyuridine; Cat. No. A10044; Invitrogen, Thermo Fisher Scientific, Waltham, MA, USA). Eyeballs were the recovered and enucleated, fixed with ROTI Histofix for 2 hours at room temperature, transferred to PBS, and kept at 4°C until dissection. Retinas were dissected out in ice-cold PBS using a stereomicroscope (Zeiss Stemi DV4 Stereo Microscope, Carl Zeiss Microscopy GmbH, Oberkochen, Germany) and being blocked overnight at 4°C in 5% donkey serum containing 0.3% Triton X-100 and 0.3% BSA in PBS.

Vascular structures and pericyte coverage were visualized by incubating retinas overnight (4°C) with anti-NG2 (rabbit, Abcam ab275024, 1:200) and anti-CD31 (rat, BD 550274, 1:100) in 1% donkey serum (0.3% Triton, 3% BSA in PBS). To identify proliferating endothelial cells, EdU detection was performed using 100 µL Click-iT reaction cocktail (2 hours, room temperature). Retinas were then incubated overnight at 4°C with anti-ERG (rabbit, Abcam ab92513, 1:100) and anti-CD31 (rat, BD 550274, 1:100) in 1% donkey serum with 0.3% Triton X-100 and 3% BSA.

Following four washes (5 minutes each) with PBS, retinas were incubated (1:200, 1 hour, room temperature) with appropriate species-specific donkey Alexa Fluor-conjugated secondary antibodies (Thermo Fisher Scientific, Waltham, United States). Samples were then mounted in Fluoromount-G mounting medium (Cat. No. 00-4958-02; Invitrogen, Thermo Fisher Scientific, Waltham, MA, USA) and covered with a glass coverslip. Images were taken with a confocal microscope (Leica SP8 or Zeiss LSM-780) and analyzed using the respective software i.e., LASX (Server version 1.9.0.3233, Leica, Wetzlar, Germany) or ZEN 2012 (blue edition) software (Zeiss, Jena, Germany) as described.^51,52^

### Cell Culture

#### Pericytes

Human brain microvascular pericytes were obtained from ScienCell Research Laboratories (Carlsbad, CA, USA) and maintained in DMEM/F-12 medium (11320033, Gibco, Waltham, MA, USA) containing 5% FCS (Gibco, Waltham, MA, USA), insulin (Sigma-Aldrich, Taufkirchen, Germany) and human FGF (Peprotech, Hamburg, Germany). Cells were used up to passage 7.

#### Endothelial cells

Human umbilical veins were obtained from local hospitals and endothelial cells were isolated and cultured as described previously,^53,54^ and used up to passage 4. Endothelial cells were cultured in ECGM2 medium (PromoCell, Heidelberg, Germany) containing 8% heat inactivated foetal bovine serum (FBS), gentamycin (25 µg/mL) non-essential amino acids (Thermo Fisher Scientific, Schwerte, Germany) and Na pyruvate (1 mmol/L, Merck, Darmstadt, Germany). EndMT was induced in some experiments by incubating cells with IL-1β and TGF-β2 (1 ng/mL, Peprotech, Hamburg, Germany) in the absence and presence of sodium lactate (10 mmol/L)for 72 hours.

#### Endothelial cell-pericyte co-culture

Pericytes were seeded (300,000 cells per well) on the underside of gelatin-coated six-well Transwell filter inserts (pore size 0.4 µm; Sarstedt, Nümbrecht, Germany). After 3 hours, the inserts were inverted and endothelial cells (600,000 cells per well) were densely plated on the upper surface. Cultures were maintained for 5 days so that endothelial cell–pericyte junctions could be established within the filter pores, as described.^55^ Cells were harvested by scraping, snap-frozen in liquid nitrogen and kept at -80°C until used. Filters were generally discarded unless they were used to assess proteins enriched in endothelial cell-pericyte junctions.

### Immunoblotting

Proteins were extracted using RIPA lysis buffer (50 mmol/L Tris/HCl, pH 7.5, 150 mmol/L NaCl, 10 mmol/L NaPi, 20 mmol/L NaF, 1% sodium deoxycholate, 1% Triton X-100, and 0.1% SDS) supplemented with protease and phosphatase inhibitors. Histones were isolated using an acid-extraction protocol. Briefly, cells were lysed in ice-cold lysis buffer containing 50 mmol/L Tris/HCl (pH 7.5), 150 mmol/L NaCl, 1% NP-40, 10 mmol/L NaPPi, 20 mmol/L NaF, 2 mmol/L sodium orthovanadate, 10 nmol/L okadaic acid, 25 mmol/L β-glycerophosphate, and 230 µmol/L PMSF. An equal amount of total protein was used for histone isolation for all samples. Lysates were centrifuged at maximum speed (≈15,000 rpm) for 15 min at 4 °C, and the supernatant was collected and stored for further analysis. Detergent-soluble proteins were resuspended in SDS-PAGE sample buffer, separated using SDS-PAGE, and processed for Western blotting as previously described.^56^

The following primary antibodies were used: anti-SLC16A3 (1:1000, Cat. 22787-1-AP, Proteintech), anti-β-actin (1:1000, Cat. A1978, Sigma), and anti-GAPDH (1:300, Millipore, Burlington, United States, MAB374), anti-acetylated-lysine (1:1000, Cat. #9441, Cell Signaling Technology; Leiden, Netherlands), anti-lactyllysine (1:1000; PTM Biolabs, Hangzhou, China; Cat. PTM-1401RM), anti-lactyl-histone H3 (Lys18; 1:1000; PTM Biolabs, Hangzhou, China; Cat. PTM-1406RM). Membranes were incubated with species-specific horseradish peroxidase–conjugated secondary antibodies diluted 1:20,000 in Tris-buffered saline–Tween. Protein bands were then visualized by enhanced chemiluminescence using a commercial ECL kit (GE Healthcare, Chicago, United States).

### RNA-sequencing

Following a five-day co-culture period of endothelial cells and pericytes, the cells were lysed on ice for 45 to 60 minutes in lysis buffer (UltraPure DNase/RNase-free water containing: 50 mmol/L Tris-HCl at pH 7.5, 150 mmol/L NaCl, 10 mmol/L MgCl₂, and 1% NP-40 and supplemented with phosphatase and protease inhibitors: 10 mmol/L NaPPi, 20 mmol/L NaF, 10 nmol/L okadaic acid, 2 mmol/L Na₃VO₄, 12 µl/ml PIM, 4 µl/ml phenylmethylsulfonyl fluoride, 200 U/ml SUPERase In RNase Inhibitor, and 25 U/ml TURBO DNaseI). To ensure complete membrane disruption, the lysate was sheared by passing it 7–10 times through a 26G needle attached to a 1 mL syringe. The homogenate was then centrifuged at 13,000 rpm for 10 minutes at 4°C. The clarified supernatant was collected and subjected to total RNA purification using the miRNeasy Micro kit, following the according to the manufacturer’s instructions.

RNA quality and library preparation integrity were assessed using a LabChip Gx Touch instrument. For library construction, 500 ng to 1 µg of normalized total RNA was processed with the SMARTer Stranded Total RNA Sample Prep Kit HI Mammalian. Sequencing was carried out on an Illumina NextSeq2000 platform with a P3 flowcell, generating 72 bp single-end reads.

Raw reads were processed with Trimmomatic (v0.39) to trim low-quality regions (quality threshold Q15 over a 5-nucleotide window) and retain reads longer than 15 nucleotides. The filtered reads were aligned to the Ensembl human genome (hg38, release 109) using STAR (v2.7.10a). Subsequent alignment filtering removed duplicate, multi-mapping, ribosomal, and mitochondrial reads with Picard tools (v3.0.0). Gene-level counts were generated with featureCounts (v2.0.4) based on exonic, strand-specific overlaps.

Differential expression analysis was performed using DESeq2 (v1.36.0) on the raw count matrix. Genes were considered significantly differentially expressed if they had an average count greater than 5, an adjusted p-value below 0.05, and an absolute log2 fold change of at least 0.585. Ensembl gene annotations were supplemented with data from UniProt.

### Mass spectrometry-based proteomics

Cell pellets were lysed in 4% SDS in 10 mM Tris-HCl pH 7.6 at room temperature, vortexed, heated at 95 °C for 5 minutes, sonicated and centrifuged. Protein lysates (50 µg) were acetone-precipitated, washed and resuspended in 6 M urea/2 M thiourea in 10 mM HEPES pH 7.4. Proteins were digested with Lys-C and trypsin at standard enzyme-to-protein ratios, and peptides were fractionated using high-pH reversed-phase columns. Peptides were analysed by LC-MS/MS using an EASY-nLC 1200 coupled to QExactive HF mass spectrometer (Thermo Fisher Scientific, Bremen, Germany) via electrospray ionization and in-house packed C18 emitters. A standard formic acid/acetonitrile gradient was used (solvent A: 0.1% formic acid; solvent B: 80% acetonitrile/0.1% formic acid). Peptide/spectrum matching, protein group assembly and LFQ were performed using MaxQuant, with full parameters deposited with the raw data. LFQ-based differential expression analysis was performed using the autonomics R pipeline (https://doi.org/doi:10.18129/B9.bioc.autonomics), which applies quantile normalization, imputation, and limma’s Bayesian moderated t-tests.

### Analysis of global acetylation and lactylation by LC-MS/MS

Whole-cell lysates were incubated with anti-acetyl-lysine (Cat. #9441, Cell Signaling Technology; Leiden, Netherlands), anti-lactyllysine (PTM Biolabs, Hangzhou, China; Cat. PTM-1401RM), or control IgG (Merck) on an end-over-end rocker overnight at 4°C. Protein complexes were captured by adding rec-Protein G–Sepharose 4B Conjugate (Invitrogen; 50 µL/mg lysate) for 2 hours, followed by centrifugation at 2200 g. Immunoprecipitates were washed with lysis buffer and then with Tris-HCl/NaCl buffer, centrifuged and stored at –20°C until processed. Pellets were resuspended in 8% sodium lauroyl sarcosinate in 100 mmol/L triethylammonium bicarbonate, boiled and reduced with dithiothreitol, followed by alkylation with iodoacetamide. Samples were diluted in triethylammonium bicarbonate, digested overnight with trypsin, acidified with trifluoroacetic acid and cleared of precipitated lauroyl sarcosinate. Peptides were purified on C18 StageTips, dried and reconstituted in 0.1% trifluoroacetic acid. Concentrations were measured and equal peptide amounts were used. For LC-MS/MS, 500 ng of peptides were loaded onto a µPAC™ Trapping column and separated on a µPAC™ HPLC column (Thermo Scientific) using a standard acetonitrile/formic acid gradient. Peptides were ionized via an Advion TriVersa NanoMate (Advion BioSciences) and analysed on an Orbitrap Eclipse Tribrid mass spectrometer (Thermo Scientific) in positive-ion mode. Data-independent acquisition was performed with MS1 resolution 60,000, followed by DIA scans across 40 windows (15 m/z). MS2 spectra were acquired at resolution 30,000.

Peptide identification and label-free quantification were carried out using DIA-NN with a library-free search (human UniProt/Swiss-Prot). Results were filtered to 0.01 precursor FDR, using deep-learning-generated *in silico* spectral libraries. Search parameters included carbamidomethylated cysteine (fixed), methionine oxidation (variable), N-terminal methionine excision, tryptic cleavage (K*, R*), up to one missed cleavage and precursor masses 300–1800 m/z (charge states 1-4). DIA-NN mass accuracy was optimized per run and matrices were filtered using run-specific protein q-value. Proteins with q < 0.01 were retained. MaxLFQ values were used for quantitation with imputation of missing values. Intensities based on a single precursor were set to NA.

### Metabolomics

Sample extraction, chromatographic separation (HILIC and C18), and LC-MS/MS quantification were performed using our previously reported workflow.^9^ Metabolites were extracted using a TFE-based protocol. Briefly, samples were treated with trifluoroethanol/water (1:1), vortexed, and incubated on ice, followed by addition of methanol/ethanol (1:1). The metabolites were extracted using a TFE-based technique. Briefly, samples were treated with trifluoroethanol/water (1:1), vortexed, and incubated on ice before being added to methanol/ethanol (1:1). Following a second incubation, ice-cold water with isotope-labeled internal standards was added. The samples were then sonicated on ice, centrifuged at 14,000 rpm for 10 minutes at 4°C, and the supernatants were collected. The extracts were divided, evaporated under nitrogen, and then freeze-dried. For HILIC-based analysis, dried materials were reconstituted in H₂O:acetonitrile (1:3), while for C18-based analysis, 0.2% formic acid was added to H₂O. Following the final centrifugation, the supernatants were transferred to MS vials for LC-MS/MS quantification.

### 13C glucose flux

Cells were cultured in previously described co-culture conditions, either as endothelial cells alone (EC-EC) or in combination with pericytes (EC-PC). Cultures were kept in their respective environments for four days. On day 5, cells were washed twice with PBS and subjected to two different experimental treatments. One set of EC-EC and EC-PC cultures were incubated in standard DMEM/F-12 medium, while the other set was incubated in glucose-free DMEM/F-12 supplemented with 2.5 mmol/L ¹³C-labeled glucose (Cat. No. 389374, Sigma-Aldrich, Taufkirchen, Germany). All cultures were incubated at 37°C for 15 minutes.

### Statistics

Results are presented as mean ± SEM. GraphPad Prism software (v. 9 to 10.4.1) was used to assess statistical significance. Differences between two groups were compared by two-tailed unpaired t-test. Differences between three or more groups were compared by one-way ANOVA followed by the Tukey’s multiple comparisons test. Experiments in which the effects of two variables were tested were analyzed by 2-way ANOVA followed by Tukey’s multiple comparisons test. Differences were considered statistically significant when P < 0.05. Only exact significant P values are reported.

## Acknowledgements

The authors are indebted to Isabel Winter, Mechtild Piepenbrock-Gyamfi and Sandrine Ngaha for expert technical assistance. This work was supported by the Deutsche Forschungsgemeinschaft (SFB 1531/1 B03 – Project ID 456687919 and CardioPulmonary Institute, EXC 2026, Project ID: 390649896).

---

**Supplementary Fig. 1.**
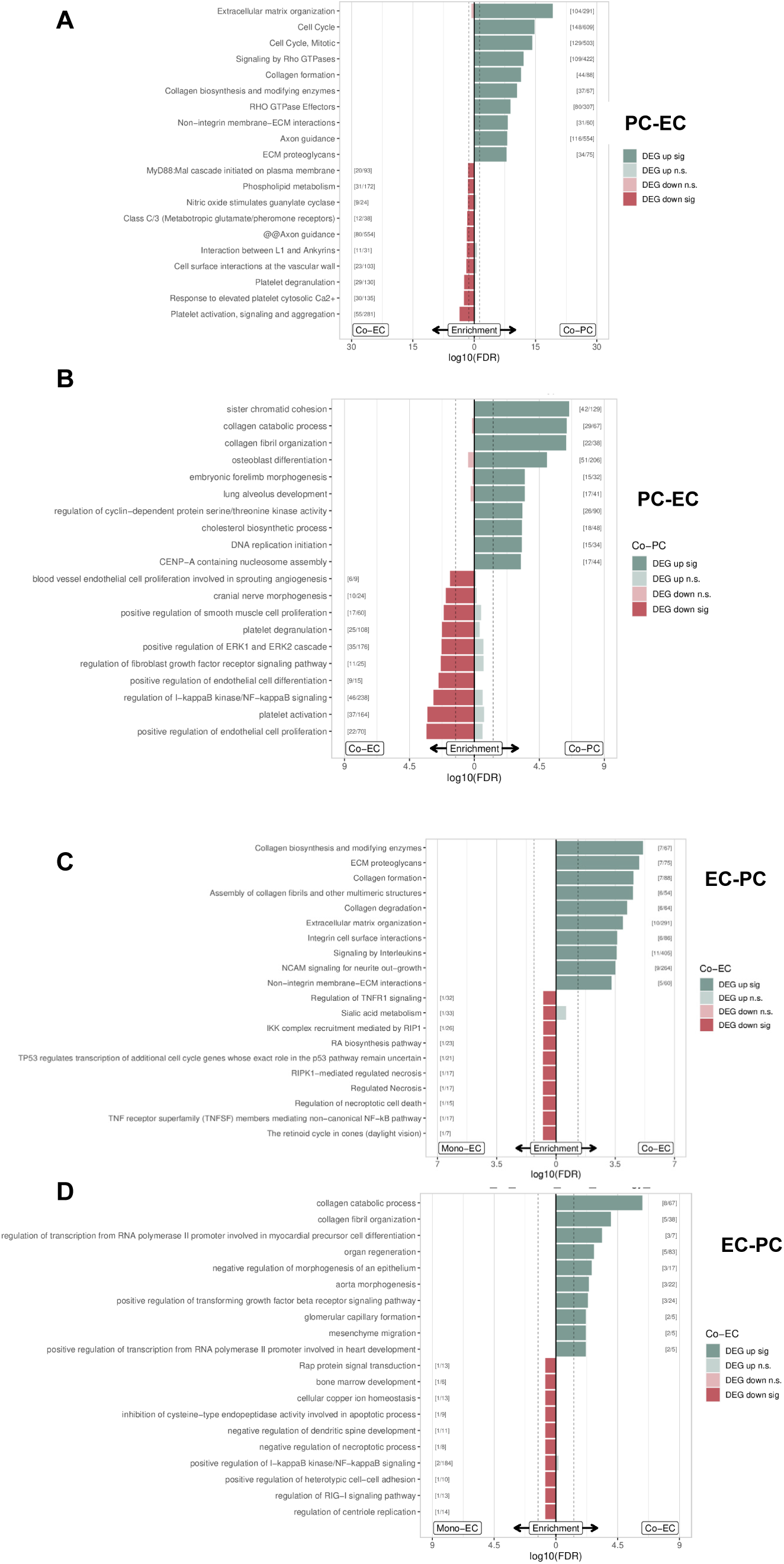
Impact of co-culture on gene expression in pericytes and endothelial cells. (**A&B**) Gene set enrichment analysis (GSEA) of the top 10 up- and down-regulated pathways in pericytes. (A) Reactome GSEA, shown are pathways significantly enriched at (FDR < 0.2). (B) Gene Ontology (GO) biological process enrichment analysis (FDR ≤ 0.2). (**C&D**) GSEA of the top 10 up- and down-regulated pathways in endothelial cells. (C) Reactome GSEA (FDR < 0.2), (D) GO biological process enrichment analysis (FDR ≤ 0.2).

**Supplementary Fig. 2.**
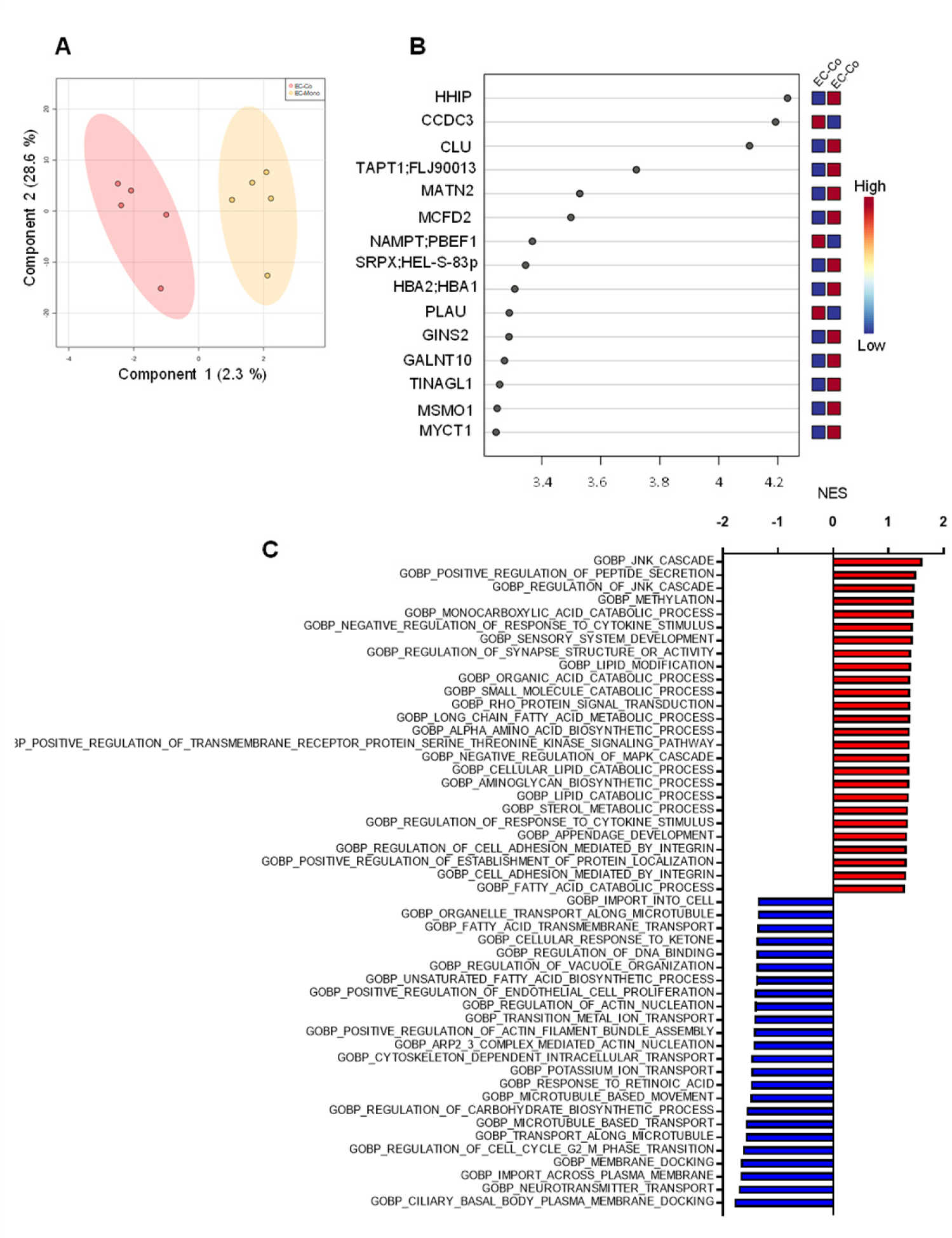
Impact of pericytes on the endothelial cell proteome. (**A**) Principal Component Analysis of components one and two in a Partial Least Squares discriminant analysis (PLS-DA), comparing the clustering of endothelial cell sin monoculture versus co-culture with pericytes. (**B**) Gene Set Enrichment Analysis (GSEA) of GO Biological Processes comparing EC/EC and EC/PC conditions. The plot shows the top enriched gene sets ranked by Normalized Enrichment Score (NES), illustrating pathways that shift significantly with the presence of pericytes. (**C**) GSEA comparing endothelial cells in mono-versus co-culture with pericytes. Red = significantly enriched in EC monoculture, blue = enriched in endothelial cells in contact with pericytes.

**Supplementary Fig. 3.**
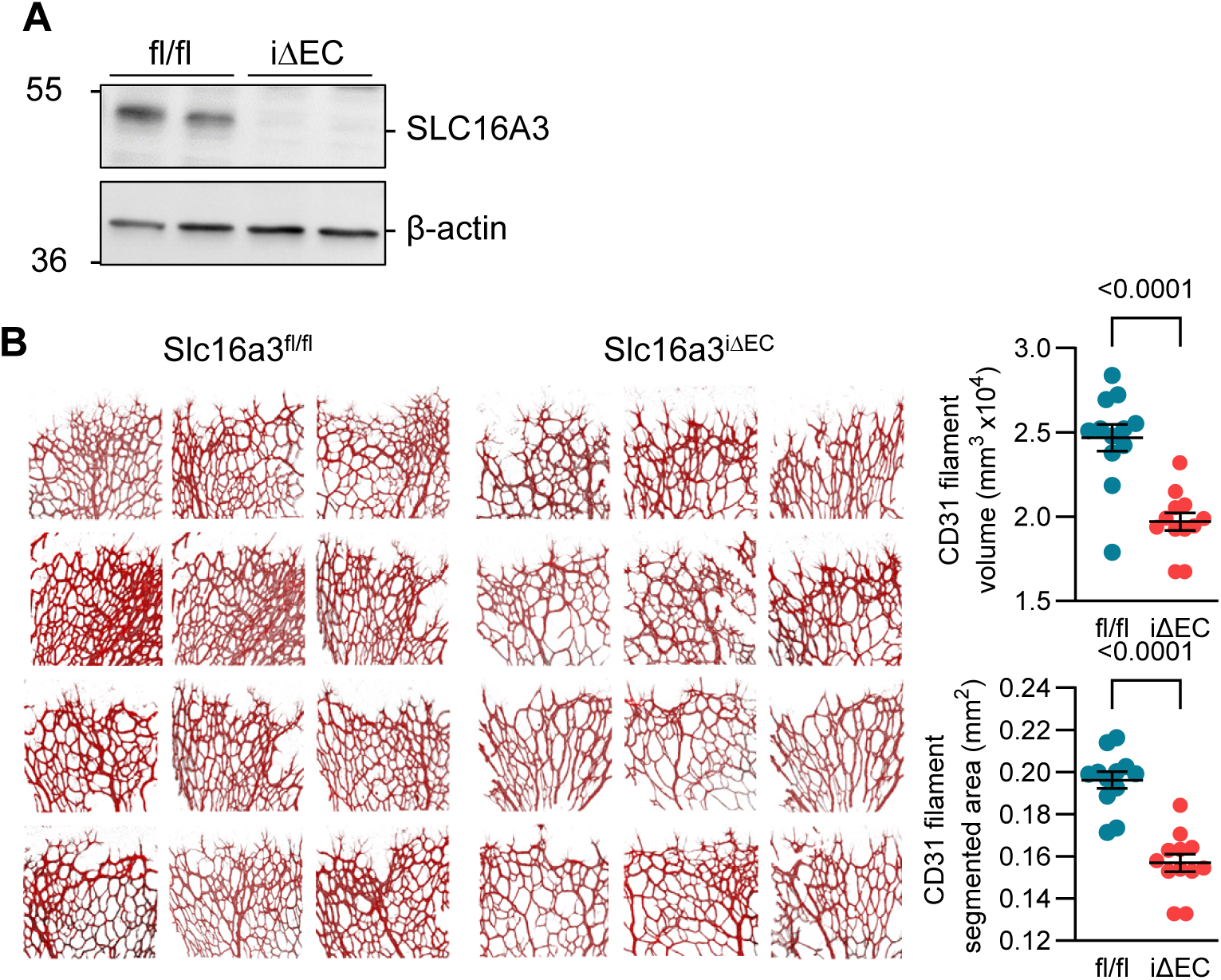
Impact of endothelial cell-specific SLC16A3 deletion on retinal angiogenesis on P6. (**A**) Western blot showing the efficient tamoxifen-induced deletion of SLC16A3 in Slc16a3^iΔEC^ mice versus Slc16a3^l/fl^ littermates. (**B**) Representative whole-mount images of the CD31+ network in retinas from Slc16a3^fl/fl^ and Slc16a3^iΔEC^ mice on P6; bar = 200 µm; n= 6 mice / group, 2 retinas / mouse (Student’s t test).

